# Modulation of cell adhesion and migration by poly-dispersed-acid-functionalized-single-walled carbon nanotubes in lung epithelial cells

**DOI:** 10.1101/2020.04.23.056895

**Authors:** Sushreesangita P. Behera, Rajiv K. Saxena

**Author notes:** **Correspondence: Prof. Rajiv K. Saxena**, Faculty of Life Sciences and Biotechnology, South Asian University, Akbar Bhawan, Chanakyapuri, New Delhi 110021.

## Abstract

Epithelial cell lining of the lung alveoli is under constant onslaught of airborne pathogens and pollutants that may cause injury and disruption of the epithelial lining. Repair mechanisms involve proliferation and migration of nearby healthy epithelial cells to the site of injury. Using murine LA4 and human A549 lung epithelial cell lines, and *in vitro* models of cell migration we have examined the modulation of cellular adhesion and migration by poly-dispersed acid-functionalized single-walled carbon nanotubes (AF-SWCNTs). Flow cytometric and confocal microscopy studies indicated that AF-SWCNTs were efficiently internalized by both cell lines and were localized essentially in the cytoplasmic area. In the scratch wound repair model, exposure to AF-SWCNTs blocked the filling of the scratched area of the cellular monolayers in both cells. Behaviour of the cells around the scratch area was examined in by live-cell imaging time-lapse micrography. The results indicated active cell proliferation around the scratch area that was totally blocked by AF-SWCNTs in LA4 cells and significantly inhibited in A549 cells. Cell migration across a porous membrane in transwell assay system also indicated a marked inhibition of migration of both cells across the membrane. Effect of AF-SWCNTs on the expression levels of important cell proteins involved in cell migration and adhesion were examined by western blotting and immunofluorescence staining. Expressions of proteins like β-Catenin, NM-Myosin and Vimentin that play crucial role in cell migration were suppressed in AF-SWCNTs-exposed cells whereas the expression levels of E-cadherin and Claudin-1, involved in cell-cell adhesion remained unaltered. Our results provide an insight into the mechanism of repair of lung epithelial cell layers.

## Introduction

Carbon nanotubes (CNTs) first reported by lijima (1991) have exceptional mechanical, thermal and electrical properties, and have found multiple commercial usage in industry as well as in biomedical fields (Chen and Yan 2017, Rezaee et al 2018, Yudasaka et al 2018, Schroeder et al 2019). The growing demand has led to an exponential increase in the production of CNTs that is about 15000 metric tons now growing at an annual rate of about 30% (nanowerk.com). Since there are no efficient mechanisms of degradation of CNTs in nature, accumulation of CNTs in environment and its adverse health effects are of widespread concern (Helland et al 2007, Poland et al 2008, Sachar and Saxena 2011, Jackson et al 2013). Pristine single walled carbon nanotubes are highly hydrophobic and exist as large agglomerates insoluble in aqueous media. CNTs can however be converted to poly-dispersed hydrophilic form by a variety of processes including acid functionalization that results in the attachment of carboxyl and sulfonate groups on carbon atoms in CNT walls (Wang et al 2006, Saxena et al 2007, Gazia and EI-Magd 2019). The resulting acid-functionalized single-walled carbon nanotubes (AF-SWCNTs) are well dispersed in aqueous media and interact well with cells.

Our group has been studying the interactions of AF-SWCNTs with cells in several biological systems, and have demonstrated modulatory effects of AF-SWCNTs on activation and functioning of T cells (Alam and Saxena 2013, Dutt and Saxena 2019a), B cells (Dutt et al 2019), antigen presenting cells (Kumari and Saxena 2012, Abbas and Saxena 2015), hematopoietic activity in bone marrow (Sachar and Saxena 2011, Bhardwaj and Saxena 2015) and lung epithelial cells (Saxena et al 2007, Kumari and Saxena 2012, Abbas and Saxena 2015). In the present study, we have focused on the modulation of adhesion and migration properties of lung epithelial cells by AF-SWCNTs, using two cell lines (LA4 murine and A549 human lung epithelial cell lines) as models. Since lungs are under constant onslaught of air-borne microbes and pollutants, that may injure the alveolar epithelial cell lining, repair mechanisms are important to maintain the integrity of alveolar epithelial lining. Injury of the lung epithelial lining triggers repair mechanisms including the migration of epithelial cells from nearby areas to the denuded surface in alveoli (Crosby and Waters 2010, Gardner et al 2010). Our results demonstrated that, AF-SWCNTs are internalized in significant amounts by the epithelial cells. Furthermore, AF-SWCNTs significantly inhibit the cell migration process in both LA4 and A549 cells by interfering with the expression of crucial marker proteins involved in the cell migration. These findings will provide essential information about the modulatory behaviour of the carbon nanotubes with respect to cellular migration of lung epithelial cells and molecules that participate in this effect.

## Materials and Methods

### Cells and Reagents

LA4 (a murine lung epithelial cell line) and A549 (a human alveolar epithelial cell line) were obtained from American Type Cell Culture, cultured in a humidified incubator at 37°C and 5% CO_2_ and maintained in RPMI 1640 culture medium supplemented with 2 mM glutamine, 4.5g glucose per litre, 10 mM HEPES buffer pH 7.2, gentamycin (10 μg/ml), and fetal bovine serum (10% V/V).

### Reagents and other supplies

Acid-functionalized single-walled carbon nanotubes prepared by arc-discharge method and carboxylated by nitric acid and sulfuric acid treatment, were procured from Sigma-Aldrich (Cat No. 652490). The material was in form of black powder that was >90% carbon and 5-8% metals with 1-3% carbon atoms as carboxyl acid groups. Average dimension of the nanotubes was 1.4 nm and its bundles ranged 4-5 nm in width and 500 to 1500 nm in length. The product was easily dispersible in water at the concentration used. Zetasizer analysis indicated a zeta potential of −45.3 mV and average size of 178 dnm (Supplementary Figure 1A). TEM picture of AF-SWCNTs indicated the fibrous structure (Supplementary Figure 1B). Mid and near infrared spectroscopic results provided by Sigma-Aldrich are given in Supplementary Figure 1C.

N-Hydroxysuccinimide (NHS), 1-ethyl-3-(3-dimethylaminopropyl) carbodiimide (EDAC), 2-(N-morpholino) ethane-sulfonic acid (MES), and Transwell 24 well plates were procured from Sigma Aldrich, USA. Alexafluor 633 hydrazide was obtained from Molecular Probes (Carlsbad, CA, USA). RPMI 1640 culture medium supplemented with glutamine (2 mM), HEPES buffer pH 7.2 (10 mM), gentamycin (10 μg/ml), and fetal bovine serum (10% V/V) was purchased from Himedia, India. Centricon 3kDa centrifugal filter device was obtained from Millipore (Billerica, MA, USA). Monoclonal antibody against NM-Myosin (ab55456) was procured from Abcam (Cambridge, United Kingdom). ß catenin (MA110056), Vimentin (OMA1-06001), E-Cadherin (PA5-16766) and Claudin-1 (PA516833) antibodies were purchased from and Thermo Fisher (Waltham, USA). Live cell imaging dish (35×10 mm) was purchased from Eppendorf (Hamburg, Germany).

### Tagging of fluorochrome to AF-SWCNTs

Fluorescence tagged AF-SWCNTs (FAF-SWCNTs) were obtained by chemically tagging the fluorochrome Alexa Fluor 633 to AF-SWCNTs by using the MES buffer containing EDAC and NHS as described elsewhere (Sachar and Saxena 2011), and were sonicated before use. The tagging efficiency of fluorophore to AF-SWCNTs was > 90% (Dutt and Saxena 2019a).

### Flow Cytometry

For studies on the uptake of FAF-SWCNTs, LA4 and A549 cells were incubated in 24 well plate [0.2×10^6^/ml/well] with or without 10 μg/ml of FAF-SWCNTs. Then cells were harvested by trypsinization, washed with PBS and analyse on FACS Verse flow cytometer (BD Bioscience).

### Confocal Microscopy

To study the uptake and localization of the FAF-SWCNTs, both the LA4 and A549 cells were cultured [0.05×10^6^/ml/well] on a coverslip in a 12 well culture plate, incubated with 10 μg/ml of the FAF-SWCNTs for 24 hours, washed twice with PBS, fixed with 4% paraformaldehyde. Nuclei were counterstained with Hoechst 33322 dye as described before (Dutt et al, 2019). Coverslips were mounted onto the glass slide with 50% glycerol and slide visualized on a Nikons A1R Confocal Laser Scanning Microscope.

### Scratch Wound Assay and Live Cell Imaging for Video Recording

To study the in vitro wound repair process, both LA4 and A549 cells were cultured with an initial seeding density of [0.5×10^6^ cells/ml/well], in a 6 well culture plate. After 24 hours, when complete monolayers were formed, AF-SWCNTs were added at concentrations of 0 (control), 20, 50 and 100 μg/ml for 6 hours. The monolayers were then scratched with the help of 200 μl pipette tip to create linear wounds. Monolayers were washed to remove dislodged cells and culture medium replaced with the respective concentrations of AF-SWCNTs. The wounds were observed and photographed at 0, 12, 24 and 36 hours by NIS-Nikons phase contrast microscope. The residual wound areas were measured using NIH Image J software. The assay was performed in triplicate and repeated thrice. For live cell imaging videography, culture dish was placed on the live imaging stage of microscope comprising transparent culture chamber maintained at 37°C and with 5% CO_2_ in air environment. Time lapse differential interference contrast (DIC) images were acquired (1 per 10 minutes) for 36 hours using confocal laser microscopy.

### Transwell assay of Cell migration

To investigate the effect of the AF-SWCNTs on cell migration, the migration ability of LA4 and A549 cells were analysed using a 24-well plate transwell system with 8-μm pore size polycarbonate filter (Catalogue no.3422, Corning, Life Science). Briefly, cells at the density of [3.0×10^4^/100 μl/well] were cultured in the upper chambers of the transwell unit in serum-free medium. The lower chambers of the transwells contained normal culture medium containing 10% FBS as chemoattractant. Cells in the upper chamber were treated with or without 50 and 100 μg/ml of AF-SWCNTs and incubation carried out at 37°C in a 5% CO_2_ atmosphere for 24 hours. The non-migrated cells on the upper side of the transwell membrane were removed using a cotton swab. Cells that migrated to the lower side of the membrane were fixed with 70% alcohol and stained with crystal violet and were visualized under Nikons phase contrast microscope. The number of migrated cells were counted with the help of NIH Image J software.

### Western Blotting

Western blot analysis was carried out as described before (Singh et al 2019) with slight modification. Briefly, LA4 and A549 cells were cultured with or without 50 and 100 μg/ml of AF-SWCNTs for 48 hours. Cells were washed twice with ice-cold PBS and incubated in Radio-immuno-precipitation assay (RIPA) lysis buffer containing a cocktail protease inhibitor mixture at 4°C for 20 min. and then stored at −80° overnight. Next day, cell lysates were collected, sonicated 3 three times for 5 sec each and analysed for protein content using the BCA protein assay kit (Thermo Fisher scientific). Samples containing 30 μl (1 μg/μl) of cell lysate proteins per lane were resolved on SDS-PAGE (8% sodium dodecyl sulfate-polyacrylamide gel electrophoresis) along with pre-stained protein ladder and transferred onto PVDF membranes (Invitrogen). The transferred membranes were blocked for 1 hour using 5% BSA in TBST (25 mm Tris-HCl, pH 7.4, 125 mm NaCl, 0.05% Tween 20) and incubated with the appropriate primary antibodies (ß catenin, NM-Myosin II, Vimentin, E-Cadherin and Claudin-1) at 4°C overnight. Membranes were washed twice with TBST for 5 min. and incubated with horseradish peroxidase-coupled isotype-specific secondary antibodies for 1 h at room temperature. The immune complexes were detected by enhanced chemi-luminescence detection system (Bio-Rad) and quantified using NIH Image J software.

### Intracellular immunofluorescence Staining

LA4 and A549 cells were cultured to confluency on glass cover slips and treated with 100 μg/ml of AF-SWCNTs for 24 hours, fixed in 4% paraformaldehyde in PBS. The samples were washed two times, permeabilized with 0.1% Triton X-100 for 15 min. and blocked with 5% bovine serum albumin in PBST for 1 hour before staining with ß catenin and NM-Myosin (1:400 dilution) primary antibody, followed by Alexa Fluor 488-conjugated secondary antibody (Thermo Fisher). The nuclei were stained with Hoechst 33342 dye and coverslips mounted with glycerol 50% for visualization using Nikons confocal laser scanning microscope.

### Statistical Analysis

All experiments were repeated at least three times. Results were expressed in Mean ±SEM. The paired t-test was performed to define the significance of all the experiments. Statistical analyses were performed using Sigma Plot software (Systat software, San Jose, CA).

## Results

### Uptake of FAF-SWCNTs by LA4 and A549 cells

The aim of this study was to examine the effect of AF-SWCNTs on cell adhesion and migration in lung epithelial cell lines. Cell lines used were LA4 (a murine lung epithelial cell line) and A549 (a human lung epithelial cell line). In order to visualize the interaction between AF-SWCNTs and the epithelial cells, the cells were incubated with fluorescence tagged AF-SWCNTs (FAF-SWCNTs) for up to 24 hours and uptake of the FAF-SWCNTs analysed by flow cytometry as well as confocal microscopy. Consolidated results of flow cytometry given in Figure 1 (panel A) show the kinetics of uptake of FAF-SWCNTs by LA4 and A549 cells and indicate that at 12 and 24 h time points almost all cells were positive for FAF-SWCNTs uptake. Mean Fluorescence Intensity (MFI) data in the same figure (right Y axis) shows that the A549 cells take up relatively more FAF-SWCNTs than the LA4 cells. For confocal microscopy, cells exposed to FAF-SWCNTs were counterstained by Hoechst 33342 dye to visualize blue stained nuclei while FAF-SWCNTs could be visualized by its red fluorescence. Results in Figure 1B (top panel) shows the red fluorescence of FAF-SWCNTs was generally localized in the cytoplasmic area of the LA4 cells. Z-sectioning images (2, 2.5 and 3 μm depths) in Figure 1B (lower panel) show that the FAF-SWCNTs were actually localized in cytoplasm with relatively poor uptake in nuclei. Similar data for A549 cells is presented in Figure 1C. These results indicate that in both the epithelial cell lines, AF-SWCNTs get rapidly internalized and accumulate predominantly in cytoplasm.

**Figure 1:**
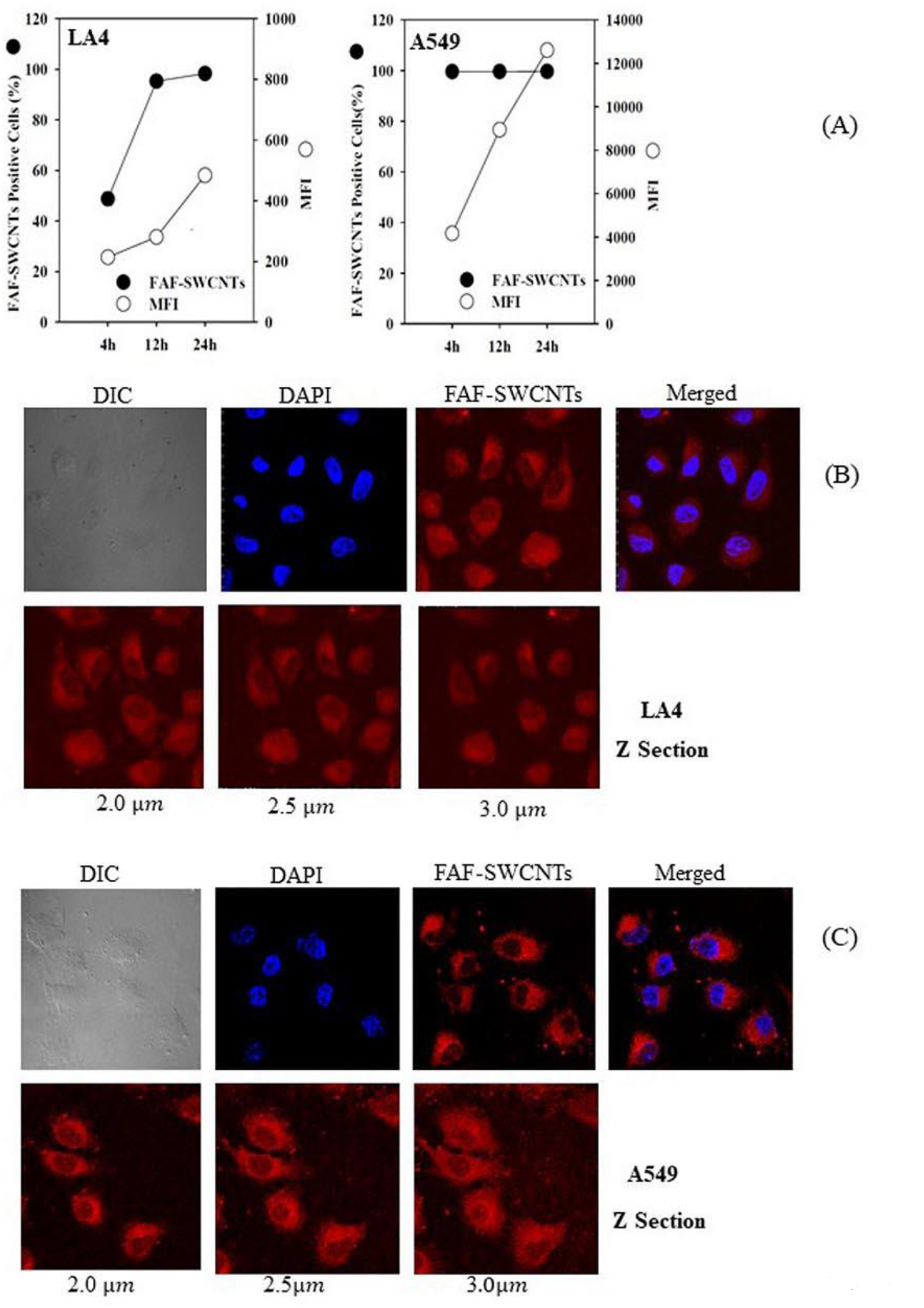
Uptake and Localization of Fluorophore-tagged-AF-SWCNTs (FAF-SWCNTs) by LA4 and A549 cells. LA4 and A549 Cells [0.2×10^6^/ml/well] were cultured with FAF-SWCNTs (10 μg/ml) for up to 24 hours. After incubation, cells were washed with PBS, detached using trypsin, and analyzed on a flow cytometer to evaluate the uptake of FAF-SWCNTs by LA4 and A549 cells. Panel A shows the kinetics of uptake of FAF-SWCNTs by both LA4 and A549 cells, where left Y-axis shows FAF-SWCNTs positive cells (%) and right Y-axis shows mean fluorescence intensity (MFI) for the uptake of FAF-SWCNTs. Each plotted value is a Mean±SEM of 3 individual experiments. Confocal microscopy results are shown in panels B (LA4 cells) and C (A549 cells). In each panel B and C, upper set of micrographs show FAF-SWCNTs uptake (red fluorescence) and nuclei are defined by blue stain of Hoechst 33342 dye, and lower set shows z-sectioning at 2, 2.5 and 3 micron depths. Magnifications 100X in all cases. DIC stands for Differential Interference Contrast.

### Inhibition of in vitro wound-healing by AF-SWCNTs

It is well known that if a scratch is made in a monolayer of epithelial cells, cells in the surrounding area move to fill the wound area (Liang et al 2007, Cory 2011, Chen 2012). Effect of AF-SWCNTs was examined in this scratch assay of wound-healing. Results in top panel of Figure 2 show that in the case of monolayers of control LA4 and A549 cells, the scratch was completely filled by 36 hours post-scratch. However, addition of AF-SWCNTs resulted in a significant inhibition of gap-filling by both cells. Kinetics data of filling of the scratch for control and AF-SWCNTs treated LA4 and A549 cells are shown in the bottom panel of Figure 2. In control cells, the scratch gap was completely filled by 36 hours but in presence of 20, 50 or 100 μg/ml of AF-SWCNTs, only 30 to 50% of the gap area could be filled in both cases. These results indicate that AF-SWCNTs reduced significantly the ability of cells to fill the scratch wound.

**Figure 2:**
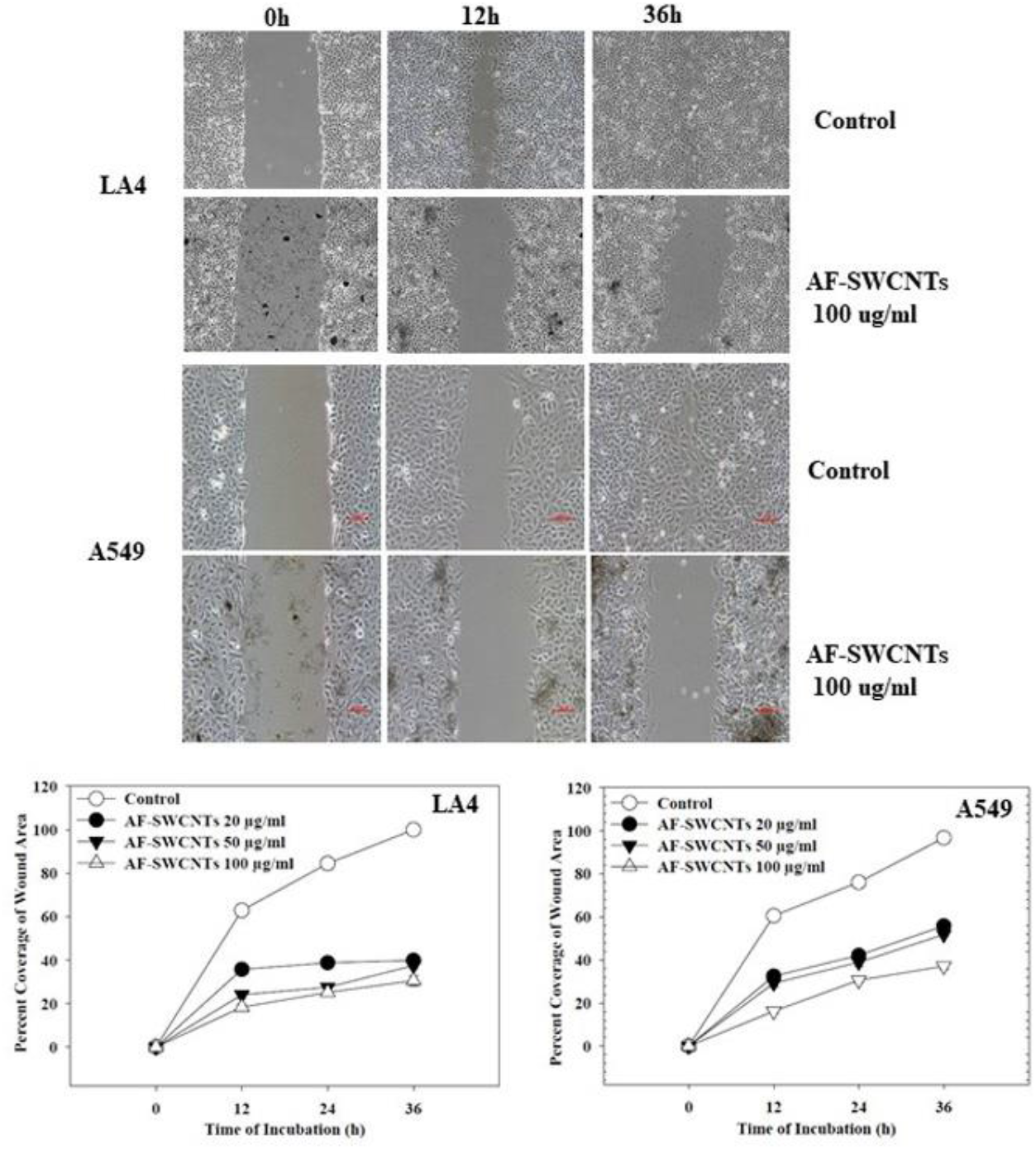
Effect of AF-SWCNTs on the In Vitro Wound Repair Process in LA4 and A549 cell monolayers. LA4 and A549 cells [0.5 ×10^6^/ml/well] were cultured in 12 well plate to get complete monolayers, after which cells were incubated with or without AF-SWCNTs [20, 50 and 100 μg/ml] for 6 hours. Uniform scratches were made in the monolayers and the wounded monolayers were washed with fresh medium and media replaced with original concentrations of AF-SWCNTs. Photographs were taken at 0, 12, 24 and 36 hours post wounding using a Nikon phase contrast microscope. Top panel shows representative images of the wound repair process in control and AF-SWCNTs (100 μg/ml) exposed LA4 and A549 cells, at 0, 12 and 36 h time points. Area of the residual scratch wounds at 0, 12, 24 and 36 h time points were measured with Image-J software and the kinetics of the filling of scratch wound has been shown in the lower panel. Each plotted value represents Mean±SEM of 9 random measurements in three individual experiments. SEM values were too small to be visible on the plot. At each concentration of AF-SWCNTs, the statistical significance of difference between the control and +AF-SWCNT curves were highly significant (p<0.001).

### Live cell imaging to study the wound healing process

Scratch wound healing in LA4 and A549 cells was also recorded on a live cell imaging system and the video clips of cellular movement provided in supplementary data (Supplementary Figure 2 (videoclips). There was a significant cell division activity around the scratch wound area that could be visually assessed by counting the number of dividing cells from the video clips of both control and AF-SWCNTs treated cells. Results in left panels of Figure 3 show that within the time period of 36 hours, 215 of the total LA4 cells and 170 of the A549 of cells in the view field divided in control plates and that number went down to about 60 each for both LA4 and A549 cells if 100 μg/ml of AF-SWCNTs were added to the culture medium. These results indicated that AF-SWCNTs suppressed the cell division activity around the scratch wound for both cell lines. Cell divisions in 5 hour time intervals during the wound-healing assay were also counted for both control and AF-SWCNTs treated culture wells. Data in the right panels of Figure 3 show that for LA4 cells, the cell divisions in AF-SWCNTs treated continued till 10 hours and then fell sharply to nil at later time periods. For A549 cells however, the cell division activity continued in AF-SWCNTs treated cells albeit at a significantly lower level than that in the control cells. Total number of cells in the view field for control and AF-SWCNTs treated LA4 and A549 cells were very similar and observed fall in cell division activity was not due to difference in total number of cells in the view fields.

**Figure 3:**
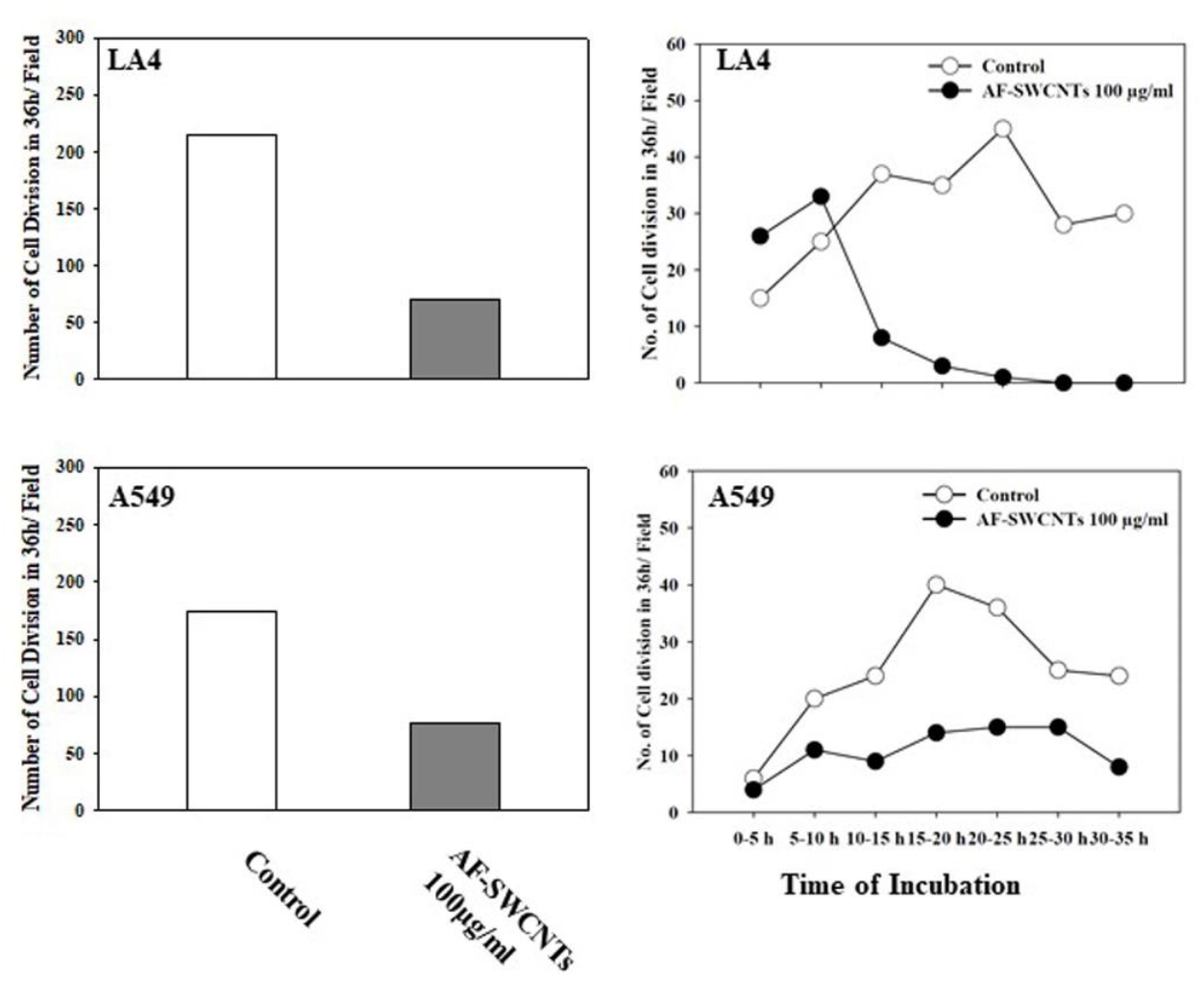
Effect of AF-SWCNTs on cellular migration and proliferation during the wound healing of scratched LA4 and A549 cell monolayers, observed by live-imaging time lapse photography. LA4 and A549 cells [0.3 ×10^6^/750 μl] were cultured in the cell imaging dish (35×10 mm). After the formation of complete monolayer, the cells were treated with 100 μg/ml of AF-SWCNTs for 6 hours, following which uniform scratches were made with the help of 20 μl pipette tip. Monolayers were washed and culture media replaced with fresh media containing the afore-mentioned concentration of AF-SWCNTs. Culture dish was placed on the stage of microscope enclosed in a small transparent culture chamber maintained at 37°C and an atmosphere of air containing 5% CO_2_. Photographs were taken at intervals of 10 minutes on Nikon confocal laser microscope (20X magnifications) for over a period of 36 hours. Actual video clips of the wound repair process in presence and absence of AF-SWCNTs for both LA4 and A549 cells has been provided in the accompanying on-line supplementary data. Actual numbers of cell divisions over the complete time span of 36 hours and within various time periods in between could be counted from the video clips. Left panels show the total number of cell division in the complete view fields in 36 hours for control and AF-SWCNTs treated cells. Right panels show the number of cell divisions during 5 hour intervals starting from 0 hours to 35 hours per field in control and AF-SWCNTs exposed LA4 and A549 cells.

### Migration of LA4 and A549 cells in trans-well assay system

Trans-well migration assay has been used by many investigators (Lohcharoenkal 2013, Chen et al 2016) to study the cellular migration from top well with serum-free medium to the bottom well containing medium with serum, through a membrane with 8 μm pores separating the two wells. Results in Figure 4 (upper panels) show the migrated cells underneath the transwell membrane, stained with crystal violet for control and AF-SWCNTs treated LA4 and A549 cells. These results show that a significantly lower number of cells migrated in if AF-SWCNTs (50 or 100 μg/ml doses) was added to the culture. Data for actual numbers of cells that migrated in control and AF-SWCNTs treated cultures is given in the lower panel of the Figure 4. These results clearly show that there was a dose dependent inhibition of cellular movement in cells exposed to AF-SWCNTs.

**Figure 4:**
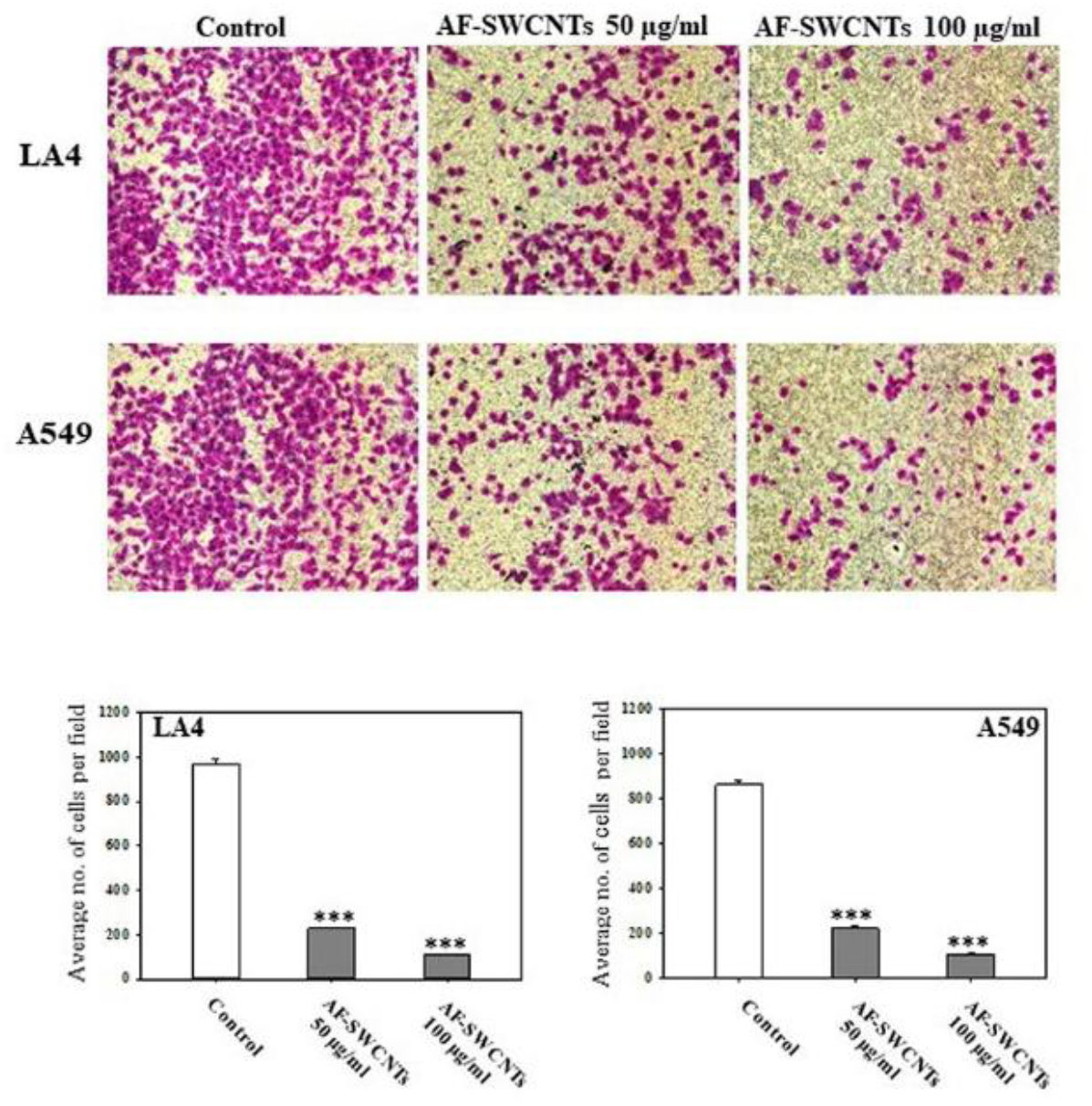
Effect of AF-SWCNTs on the cell migration of LA4 and A549 cells through a porous membrane using Transwell assay plates. LA4 and A549 cells [3 ×10^4^/ml/well] were cultured in the upper chamber of the 24 well transwell plates, containing plain serum free media. Lower chambers were filled with culture medium containing 10% FBS. After 10 min. of incubation, 50 and 100 μg/ml of AF-SWCNTs were added in the upper chambers and incubation continued for 24 hours. Transwell membranes were fixed and stained with crystal violet. The cells which migrated to the lower side of the upper chamber were visualized by Nikon Phase Contrast Microscope (10X Magnification). Top panel Images show the migration of control cells and cells treated with 50 or 100 μg/ml of AF-SWCNTs. Bottom panel shows the quantitative evaluation of average number of cells migrated per field in control and AF-SWCNTs exposed LA4 and A549 cells. Each bar represent mean ± SEM of 9 random fields in three individual experiments. Inhibition of migration in AF-SWCNTs exposed cells was highly significant (***p<0.001).

### Modulation of expression of molecules involves in cellular adhesion and migration by AF-SWCNTs

As a significant inhibitory effect of AF-SWCNTs was seen on the migration of LA4 and A549 cells in the wound-healing and trans-well migration assays, we investigated the expression levels of a some selected crucial proteins known to be involved in regulation of cell migration and adhesion, namely ß catenin, Non Muscle Myosin II (NM-Myosin II), Vimentin, E Cadherin and Claudin-1. Western blotting results in Figure 5 show that ß catenin was significantly downregulated in LA4 and A549 cells exposed to 100 μg/ml of AF-SWCNTs; the decline being 98% and 79% in LA4 and A549 respectively (quantitative band intensity in first from top panel). Expression of other crucial proteins like NM-Myosin II and Vimentin were like-wise affected by AF-SWCNTs exposure. LA4 cells, treated with AF-SWCNTs (100 μg/ml) the expression of NM-Myosin and Vimentin were 63% and 58% downregulated, while in A549 cells, the expression of NM-Myosin and Vimentin were found to be 76% and 56% downregulated in comparison to the control cells. The expression of cell adhesion molecules like E-Cadherin and Claudin-1 were not significantly altered in AF-SWCNTs treated LA4 and A549 cells.

**Figure 5:**
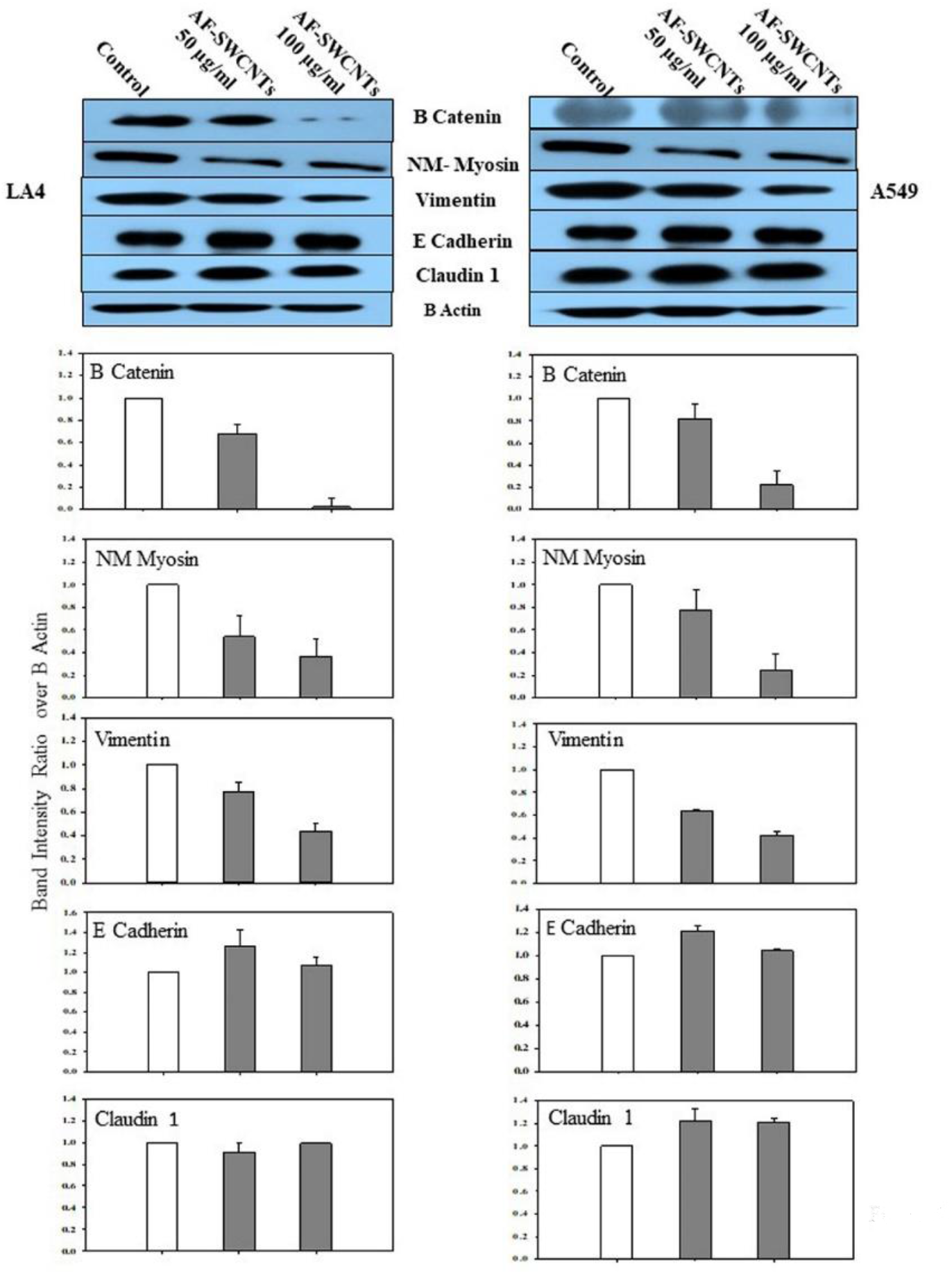
Expression of key protein markers involved in cellular adhesion and migration in control and AF-SWCNTs treated LA4 and A549 cells by Western blotting. LA4 and A549 cells [0.5 × 10^6^/ml/well] were seeded in wells of 6 well plate with or without AF-SWCNTs [50 or 100 μg/ml] for 48 hrs. Protein lysates were prepared by using RIPA extraction buffer. The estimation of protein was done by BCA method. The samples were resolved with 8% SDS-PAGE, transferred onto PVDF membrane, nonspecific binding was blocked with 5% BSA and then probed with ß catenin, NM-Myosin II, Vimentin, E-Cadherin and Claudin-1 overnight at 4 degree Celsius. The immune-complexes were detected using HRP conjugated secondary antibodies and quantified by Image-J software. Histograms show result of 3 independent experiment (Mean + SEM). Differences are significant from control with *p<0.05, **p<0.01, ***p<0.001.

Expression of β-Catenin and NM-Myosin in control and AF-SWCNTs exposed cells was further examined by immunofluorescence microscopy. For this purpose, cells were cultured with or without 100 μg/ml of AF-SWCNTs for 24 hours, fixed with 4% paraformaldehyde and permeabilized with 0.1% triton-X 100 followed by staining with β-Catenin and NM-Myosin II antibodies followed by FITC tagged secondary antibody. The expression of the two markers was examined by confocal laser microscopy. Results in Figure 6A and 6B show that AF-SWCNTs exposed cells showed decreased fluorescence intensity of both markers over control cells (Figure 6A for Β-Catenin expression, Figure 6B for NM-Myosin expression). Quantitative analysis of mean fluorescence intensity plot (Figure 6C) indicates significantly lower fluorescence intensity values as compared to the control cells in case of both LA4 and A549 cells. These results confirmed the Western blot results for these two markers.

**Figure 6:**
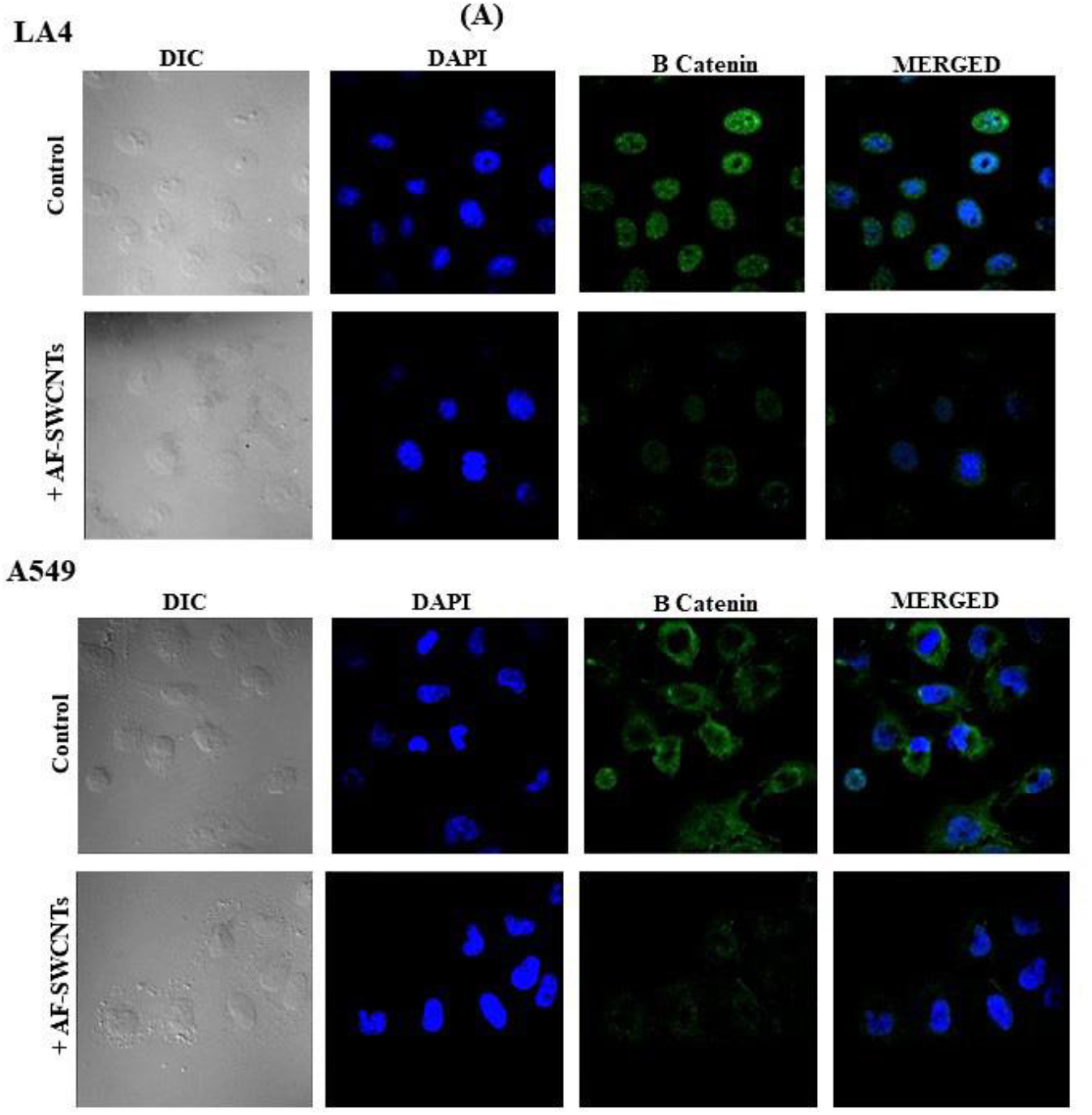
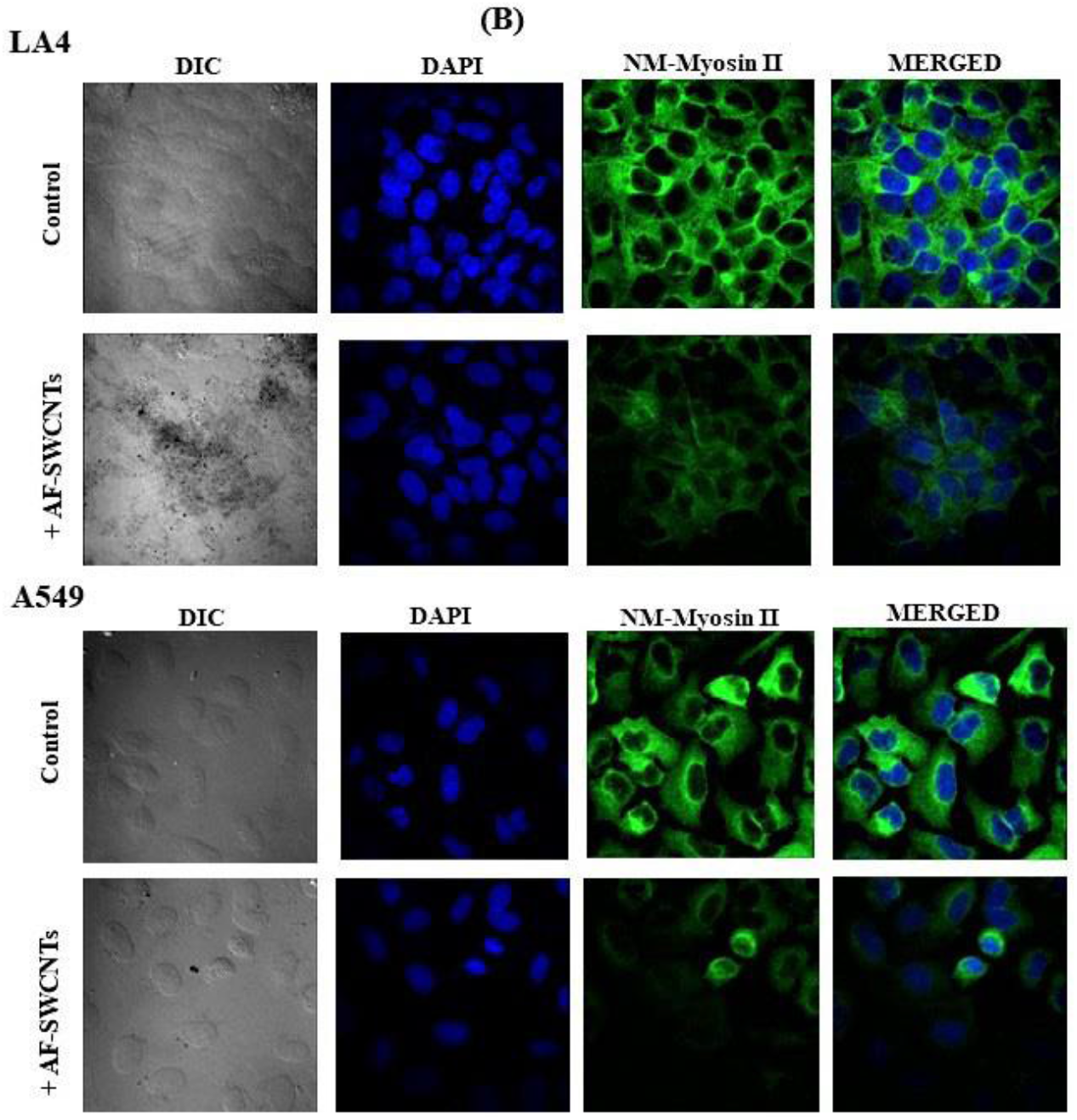
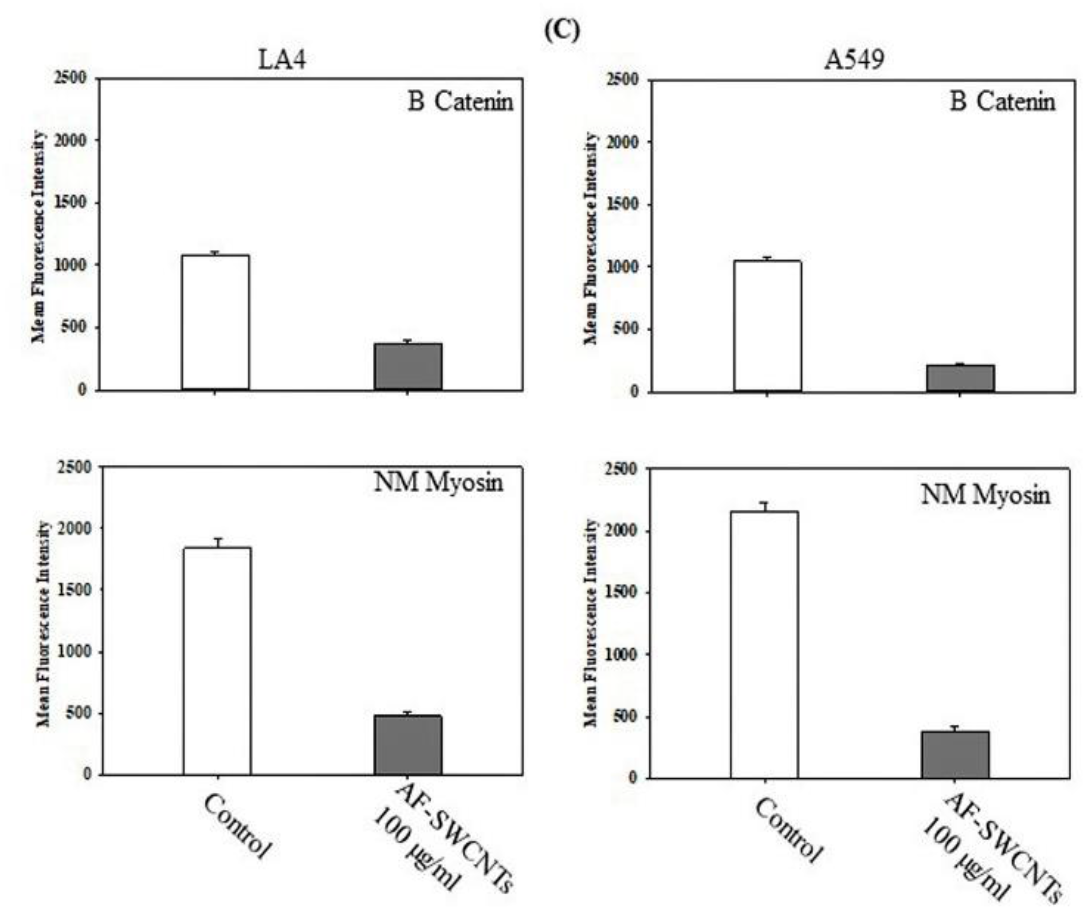
Effect of AF-SWCNTs on the expression of ß catenin and Non Muscle-Myosin in LA4 and A549 cells by Immunofluorescence assay. LA4 and A549 cells [0.5 × 10^6^/ml/well] were cultured on glass coverslip in 24 well culture plate, and treated with 100 μg/ml of AF-SWCNTs for 24 hours. Cells were fixed with 4% paraformaldehyde, permeabilized with 0.1% triton X-100 and nonspecific binding blocked with 5% BSA. The samples were incubated with primary ß catenin or NM-Myosin antibodies overnight at 4 degree Celsius, and probed with fluorescence tagged secondary antibody for 1 hour at room temperature. Hoechst 33342 dye was used to stain nucleus. The coverslips were mounted using 50% glycerin and visualized using Nikon AIR confocal laser scanning microscope. Results in Figure 6A show the distribution of ß catenin stain in LA4 cells with or without AF-SWCNTs (top two panels) and similar data for A549 cells (bottom two panels). Figure 6B show the same data for NM Myosin stain. All microscopic conditions were set the same between samples for fluorescence intensity comparison using Nikon AIR analysis software. Mean fluorescence intensity (MFI) was calculated from 8 different portion of slides in an automated manner that considers the intensity per cells in each portion of slides. Figure 6C shows quantitative differences in the MFI of control and AF-SWCNTs treated LA4 and A549 cells (mean ±SEM). ***p<0.001.

## Discussion

Cell adhesion and migration seen from the perspective of lung physiology, are crucial to sustain the integrity of lung alveoli and their repair after injury. Alveoli are lined by lung type I and II epithelial cells and cell to cell adhesion in between the epithelial cells results in an epithelial cell lining through which gaseous exchanges take place. Alveolar surface is also under constant onslaught of airborne pathogens and pollutants that may often cause localized injury and disruption of the epithelial cell surface lining the alveoli. Efficient mechanisms are in place to ensure that following injuries the epithelial cell layers get repaired quickly to permit uninterrupted life sustaining exchanges of oxygen and carbon dioxide. It is believed that epithelial cells in the vicinity divide and migrate to the site of injury to fill the gap in epithelial cell layers (Manicone 2009, Crosby and Waters 2010, Ghose et al 2017, Olajuyin et al 2019). This repair response requires that the epithelial cells participating in the repair mechanism divide and migrate to the site of injury. Factors that alter the adhesion and migratory functions of the epithelial cells may therefore influence the repair process. In the present study, we have focused upon the modulation of cellular adhesion and migration of lung epithelial cells by carbon nanotubes. It is important to examine the effects of carbon nanotubes since environmental and occupational exposure to carbon nanotubes may occur in view of the huge quantities of CNTs that are currently being manufactured for various commercial and biomedical applications. There are no efficient degradation mechanisms for CNTs in nature and their accumulation in environment and human exposure is therefore imminent.

Since highly hydrophobic CNTs exist as huge insoluble agglomerates in aqueous media, a poly-dispersed acid-functionalized form of single-walled carbon nanotubes (AF-SWCNTs) was used in our study and their effects examined on two lung epithelial cell lines [a mouse (LA4) and a human (A549)], as both these cell lines have extensively been used as models for studying the lung epithelial cells function (Sonar et al 2010, Ren et al 2016). While both cell lines efficiently internalized AF-SWCNTs and showed 100% positivity for AF-SWCNTs after 12 hours of incubation, A549 accumulated more AF-SWCNTs as indicated by significantly higher MFI values (Figure 1). Confocal laser microscopy and the Z-sectioning studies reveal that, most of the internalized carbon nanotubes were localized in the cytoplasmic space while a much smaller number transgressed nuclear membrane and was visible inside the nucleus. The mechanism of uptake of carbon nanotubes is complex and many uptake mechanisms like receptor-mediated or non-receptor mediated endocytosis, diffusion, membrane fusion, or direct pore transport of the extracellular material into the cell may contribute to this process. Several parameters such as the size, length, nature of functional groups, hydrophobicity and surface chemistry of CNTs plays crucial role in the process of internalization into the cells (Oberdorster et al. 1996, Veronesi et al 2002, Lee and Geckeler 2010, Nagai and Toyokuni 2012, Lacerda et al. 2012, Rastogi et al. 2014). The process seems to be partially energy dependent as the AF-SWCNTs uptake is lower at 4°C as compared to 37°C (Dutt and Saxena, 2019). The piercing nature resulting from its needle like structure may also be a determining factor for the nanotubes to enter into the cell through the cell membrane (Shvedova et al. 2012, Nagai and Toyokuni 2012, Lacerda et al. 2012, Dutt et al 2019). Once inside the cells, AF-SWCNTs block the ability of both cells to repair scratch wounds (Figure 2). We have used the live-cell imaging system to obtain time lapse video clips of the two cells moving in to fill the void created by scratch. Time lapse videography brings into focus two interesting features of wound healing in this system. Firstly, (a) significant cell division activity was observed around the area of scratch, and (b) migration of cells from the edge towards the scratch gap was clearly seen for both cells. Kinetics of change in cell proliferation activity recorded in the video clips show that while in the case of control cells, cell division activity continued throughout, in AF-SWCNTs treated LA4 cells, the proliferative activity ceased all together after 15 hours. In A549 cells the proliferation continued albeit at a significantly lower rate (Figure 3). In LA4 cells, the filling of the scratch gap also stopped after 12 hours time-point that correlated with the stoppage of cell division activity (Figure 3 upper panel). Migration of cells was significantly suppressed in A549 cells also but interestingly, some filling of gap appeared to have continued after 12 hour time point as suggested by the slopes of the wound space filling kinetics curves between 12 h to 36 h time points (Figure 3 lower panel). It is possible that continued cell division even though at lesser rate, could be a factor in gap filling after 12 hours in A549 cells. In the scratch wound healing system both cell division as well as cell migration could be factors in the filling of the scratch gap. In another system of cell migration across trans-well membrane potent inhibitory effect of AF-SWCNTs could be seen on cellular migration. Cell division activity may not be a prominent factor determining migration in the transwell assay and these results clearly indicate that cell migration of both cells was blocked by exposure to AF-SWCNTs.

It can be conceived that in lung epithelial cell layer repair *in vivo* as well as in scratch wound repair assay *in vitro*, moving cells would have to get detached from surrounding cells of the monolayer and initiate the process of migration. It was therefore important to assess the expression levels of molecules that participate in cell adhesion and migration and determine the effect of AF-SWCNTs on the expression of these molecules. We selected five crucial proteins β-Catenin, NM-Myosin II, Vimentin, E-cadherin and Claudin-1, which play important role and are associated with the cell migration process (V-Manzanares et al. 2009, Devreotes and Horwitz 2015, Yang et al. 2017). Significant decrease in the expression of β-Catenin, NM-Myosin II and Vimentin were observed in both LA4 and A549 cells exposed to AF-SWCNTs with respect to the control cells. β catenin is the master regulator of Wnt signalling pathway that drives the expression of genes necessary for the epithelial to mesenchymal transition associated with cell motility. (Muller et al. 2002, Yang et al. 2017). For the cell movement, the contraction force is provided by NM-Myosin II protein at the leading edge that coordinates with actin polymerization and is essential for the wound closure. The contractility of myosin also plays a strong role in directional migration and polarization of migration machinery. (Devreotes and Horwitz 2015, Tamada et al. 2007, V-Manzanares et al. 2009). Vimentin promotes the cell migration process by guiding the growth of microtubule plus end in epithelial cells. In addition to increasing cell motility, it also induces physical changes in cell shape, loss of cell–cell contacts, and increased turnover of focal adhesions, factors conducive for migration and metastasis of cancer cells. Integration of mechanical input from the environment and the dynamics of microtubules and the actomyosin network play crucial role in vimentin promoted cell migration process (Mendez et al. 2010, Rogel et al. 2011, Battaglia 2018). In AF-SWCNTs exposed LA4 and A549 cells, we found no significant changes in the expression of E Cadherin and Claudin −1 with respect to the control that is expected when cell migration is inhibited. E-cadherin is a key molecule of adherens junctions that serves as bridges connecting the cytoskeleton of neighbouring cells through direct interaction. Numerous studies have suggested a tumor suppressor role of E-Cadherin and Claudin-1 as downregulation of their expression occurs in invasive cancer cells (Gottardi et al 2001, Pećina-Slaus N 2003, Zhou et al 2015). E-cadherin and Claudin-1 are important components of the tight junctions responsible for cell– cell adhesion (Frixen et al. 1991, Zhai et al. 2008, Zhou et al. 2015). Immunofluorescence studies indicated a sharp decline in the expression of ß catenin and NM-Myosin-II in AF-SWCNTs exposed cells. This finding confirmed the result obtained by the Western blotting protein analysis.

In summary, we have demonstrated that the exposure of lung epithelial cells to AF-SWCNTs significantly inhibit the wound healing and cell migration process in LA4 and A549 lung epithelial cells. Expressions of several proteins like ß -Catenin, NM-Myosin and Vimentin, that play crucial role in cell migration were suppressed in AF-SWCNTs exposed cells whereas the expression levels of E-cadherin and Claudin-1, involved in cell-cell adhesion remained unaltered. Our results thus provide an insight into the mechanism of repair of lung epithelial cell layers.

## Acknowledgement

Research funding from the Department of Science and Technology, Government of India, and fellowship support to SPB from ICMR are gratefully acknowledged.

---

**Supplementary Figure 1.**
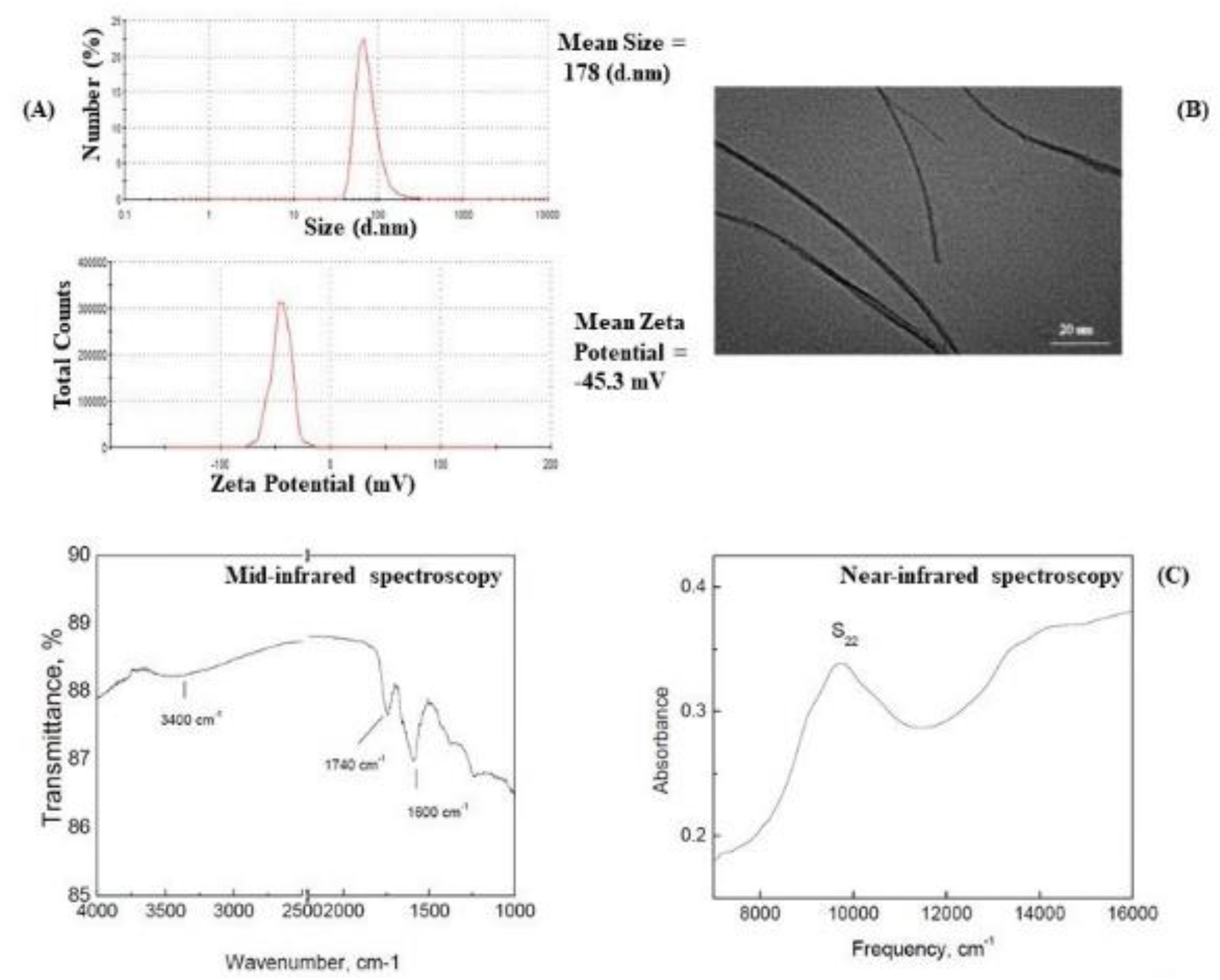
Characterization of AF-SWCNTs: Panel A shows the Zeta potential and Zeta size of AF-SWCNTs (Our data using Zeta Sizer). Panel B shows the TEM image of AF-SWCNTs (Our data). Panel C shows the mid- and near-infrared spectroscopic results (Data from Sigma-Aldrich).

**Supplementary Figure 2 (Video clips)**.

**Live cell imaging to study the in vitro wound healing process**.

LA4 and A549 cells [0.3 ×10^6^/750 μl] were cultured in the live cell imaging dish. After the formation of complete monolayer, the cells were treated with 100 μg/ml of AF-SWCNTs for 6 hours, following which uniform scratches were made in monolayers with the help of 20 μl tip. Monolayers were washed and original culture media (control or +AF-SWCNTs) was replaced in each case. Wound repair activity was monitored in a live imaging system and time lapse images were recorded (1 per 10 minutes for 36 hours). Video clips are numbered as follows: Video Clip 1: control LA4 cells; Video Clip 2: LA4 cells + AF-SWCNTs; Video Clip 3: control A549 cells; Video Clip 4: A549 cells + AF-SWCNTs. In Video Clip 2, red arrow shows a cell in LA4 cells treated with AF-SWCNTs that tries to divide but daughter cells fails to separate. A similar occurrence has also been pointed out in Video Clip 4.

